# Collagen constitutes about twelve percent in females and seventeen percent in males of the total protein in mice

**DOI:** 10.1101/2022.11.21.517313

**Authors:** Katharina Tarnutzer, Devanarayanan Siva Sankar, Joern Dengjel, Collin Y. Ewald

## Abstract

Collagen has been postulated to be the most abundant protein in our body, making up one-third of the total protein content in mammals. However, to the best of our knowledge, a direct assessment of the total collagen levels of an entire mammal to confirm this estimate is missing. Here we measured hydroxyproline levels as a proxy for collagen content together with total protein levels of entire mice or of individual tissues. Collagen content normalized to the total protein is approximately 0.1% in the brain and liver, 1% in the heart and kidney, 4% in the muscle and lung, 6% in the colon, 20-40% in the skin, 25-35% in bones, and 40-50% in tendons of wild-type (CD1 and CB57BL/6) mice, consistent with previous reports. Mice consist of 37 mg of collagen and 265 mg of protein per g of body weight. To our surprise, we find that collagen is approximately 12% in females and 17% in males of the total protein content of entire wild-type (CD1 and CB57BL/6) mice. High-Performance Liquid Chromatography approaches confirmed a 10-12% collagen over total protein estimates for female mice. Collagen staining methods and extracellular matrix-enriched proteomics estimated 5-6% of collagens over the total protein extracted. Although collagen type I is the most abundant collagen, the most abundant proteins are albumin, hemoglobulin, histones, actin, serpina, and then collagen type I. Analyzing amino acid compositions of mice revealed glycine as the most abundant amino acid. Thus, we provide reference points for collagen, matrisome, protein, and amino acid composition of healthy wild-type mice that are important for tissue and biomaterial engineering and for the comparison of these factors in various disease models.

## Introduction

Progress in biomedical science depends on building models using existing data to predict the outcome of a future experiment and either confirm or adapt the working model ^1^. These models then should be derived from data and not from assumptions. One such assumption used as a point of reference is the statement: “collagen is the most abundant protein making up one-third or 30% of total protein in mammals or vertebrates.” We found over 50 publications making such a statement without any reference or citing a review paper. It seems as if this statement is a given textbook fact (BNID 109731) ^2^ (https://en.wikipedia.org/wiki/Collagen, accessed 05.10.22); however, the actual quantifications are elusive. The biochemical properties of collagens have been studied since the end of the 18th hundred ^3^. Given the recent increased interest in collagen research of over 8000 papers published per year (Fig. 1a, Supplementary Table 1), it might be essential to have a reference point for collagen levels in tissues and entire organisms. For instance, quantification of collagen levels is important in different disease settings, such as fibrosis (too much collagen deposition ^4^ and 45% of all human deaths ^5^), immune-induced degradation of collagen in arthritis ^6,7^), more collagen and stiffer tumor environment ^8,9^, excessive collagen in skin diseases (scleroderma, keloids) ^10,11^, cancer patient survival correlates with specific collagen types ^12^, fibrotic collagen remodeling in cardiovascular diseases ^13,14^, and aging usually associated with lower collagen levels ^15^.

**Fig. 1.**
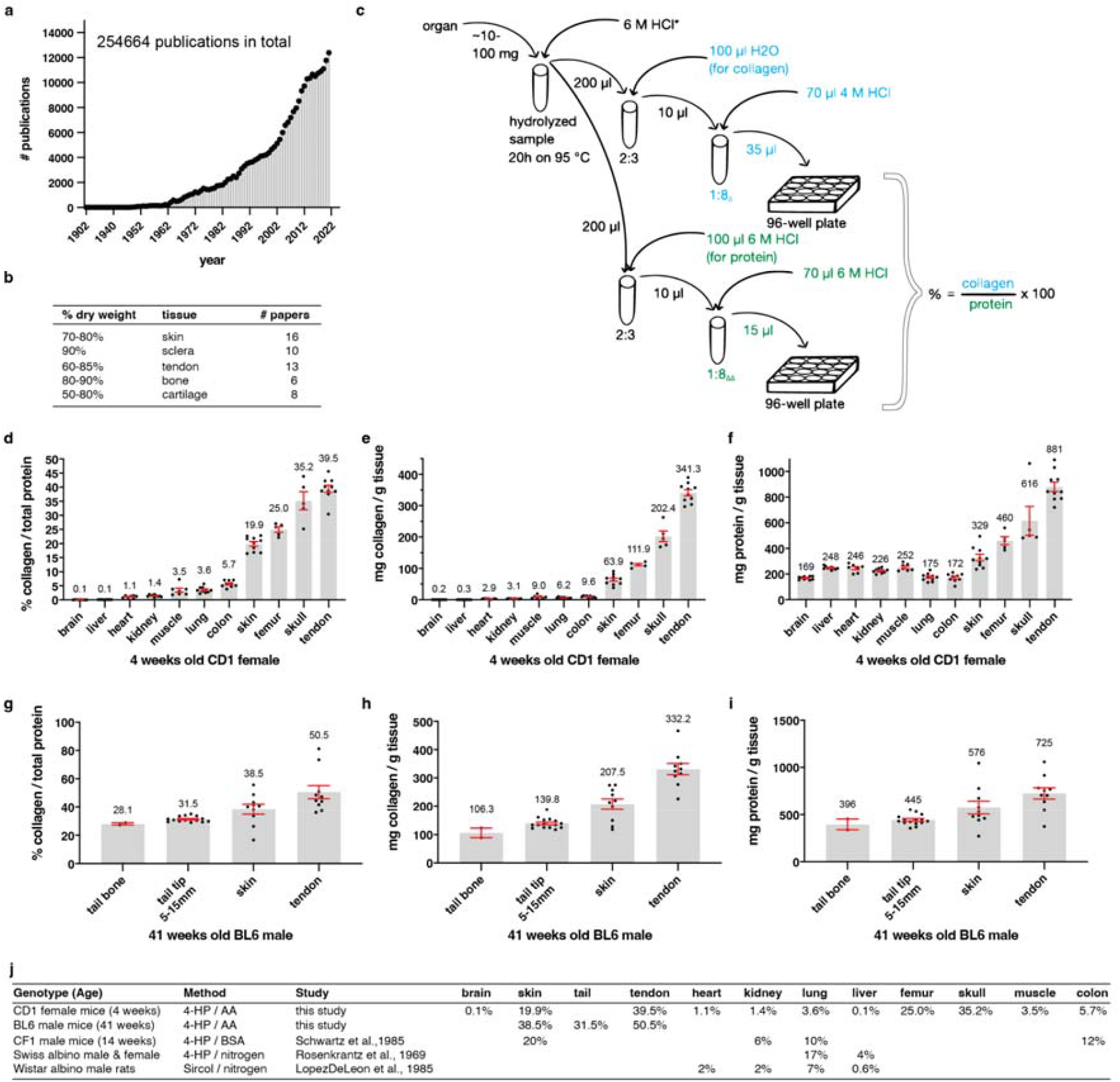
Relative collagen levels in organs. **a**, number of publications per year about collagen. See Supplementary Table 1 for details. **b**, previous studies stating the percentage of collagen in organs normalized to dry weight. See Supplementary Table 2 for details. **c**, schematic representation of the collagen and protein quantification assay for organs. * indicates dilution calculation for organs: v=aw x D_organ_ -aw (v = μL 6M HCl added to organs, aw = organ assay weight (mg), D_organ_= experimentally determined organ specific dilution factor). Δ indicates that liver and brain collagen samples were diluted 1:1 and tendon collagen samples 1:16. ΔΔ indicates that liver and brain protein samples were diluted 1:10. **d** and **g**, the percentage of collagen to the total protein of different CD1 female and C57BL/6 male organs, respectively, calculated as shown in (c). CD1 tissue is derived from 4 weeks old female mice, and C57BL/6 organs were withdrawn from 41 weeks old male mice. **e** and **h**, collagen values normalized to wet weight (mg/g). **f** and **i**, protein content normalized to wet weight (mg/g). **d-i**, shown are means ± SEM of 2-15 biological replicates (represented by dots). Each biological replicate value was derived from 1-3 technical replicates, measured in duplicates. See Supplementary Tables 4 and 5 for details. **j**, an overview of collagen content in different mouse strains and species. See Supplementary Table 5 for details.

Collagen is a fascinating biomaterial that is highly biocompatible and used for many biomedical applications in processes, such as wound healing, tissue repair, and dental implants ^16^. Collagen-based materials have been used for medical applications already reported by Galan two millennia ago for reconstructive surgery on tendons and wound closure on gladiators ^17^. Collagen can survive longer than other biomolecules, such that collagen has been extracted from dinosaur bones ^18^, and is used for estimating the age of a specimen ^19,20^. During evolution, the invention of collagen probably enabled the transitions from unicellular to multicellular organisms solving three major problems, cell and tissue adhesion, mechanical support (skeleton), and protection in the form of barrier function and mechanical shock absorption ^21^. Collagen is characterized by its triple helix repeats of (Gly-X-Y)n whereby Gly stands for glycine, X and Y can be any amino acid but most commonly are proline and hydroxyproline (reviewed in ^22,23^). For sterical reasons, the glycine on every third position is essential for adequately forming the triple helix composed of the three collagen chains (reviewed in ^22,23^). Hydroxyproline is relatively unique to collagens (only in a few other proteins found, *e*.*g*., elastin, argonaute-2, HIF-1α) ^24^ and is essential for stabilizing the collagen triple helix (reviewed in ^22,23^).

There are several methods for the detection of collagen levels: quantifying hydroxyproline levels, collagen stainings, densitometry of collagen bands in sodium dodecyl sulfate-polyacrylamide gel electrophoresis (SDS-PAGE), enzyme-linked immunosorbent assay (ELISA) coupled with collagen antibodies, immunohistochemistry using collagen antibodies, quantitative mass spectrometry enriched for extracellular matrix proteins ^4,25–29^. The gold standard in the field for assessing collagen levels is via hydroxyproline quantification ^25,29^.

Here, we hydrolyzed entire mouse samples to amino acid levels and used hydroxyproline and amino acid levels as a proxy to estimate the total collagen over total protein levels. We find two times lower total collagen levels than previously claimed in the literature. Our proteomics data suggest that collagen might not be the most abundant protein.

## Results

### Quantifying collagen levels of tissues

The importance of collagens for engineering, biological, and medical research is reflected in the exponential increase in publications about collagens over the last 60 years (Fig. 1a, Supplementary Table 1). In the literature, it has been stated that collagen makes up about 60-90% of the dry weight of bone, skin, sclera, and tendon (Fig. 1b). These percentages seem to be a given in the field since many of these statements have either no reference or cite review publications (Fig. 1b, Supplementary Table 2). Our literature search to identify the source for these percentages led to a review paper by Grant from 1972 ^30^ that summarized the collagen content of various tissues and animals. For the original work, we found two publications by Lowry and colleagues from 1941 ^31^ and Neuman and Logan 1950 ^32^ that measured the collagen content of dried tissue. However, we found no current experimental or quantitative evidence. To fill this gap, we measured total collagen levels in mice. Instead of normalizing to the organs’ dry weight, we decided to normalize to total protein levels. We harvested mouse tissues and hydrolyzed them with hydrogen chloride to the amino acid levels (Fig. 1c, Materials and Methods for details). And then e made aliquots of these amino acid homogenates to quantify total collagen and protein levels. A colorimetric read-out was used to determine free hydroxyproline amino acids that were oxidized with chloramine-T to pyrrole and stained with Ehrlich’s reagent ^33^. We normalized these hydroxyproline amino acid levels to a standard curve of rat tail collagen to determine total collagen levels (Fig. 1c). We validated our hydroxyproline and collagen quantification with an alternative but comparable protocol approach (Supplementary Fig. 1 and Supplementary Fig. 2 a-e). Since standard protein quantification does not work with free amino acids in an acidic environment, we used the color change of genipin when it reacts with amino acids as a read-out ^34^. We normalized these amino acid levels to a standardized curve of bovine serum albumin protein (BSA) to determine total protein levels (Fig. 1c; Materials and Methods for details). Thus, with this protocol, we overcame the obstacle of the insolubility of collagens in the extracellular matrix and were able to quantify total collagen over total protein (Supplementary Fig. 2f).

### Collagen levels across mice tissues range from 0.1 to 51% of total protein levels

We harvested organs from four weeks old outbred wild-type CD1 female mice and 41 weeks old inbred wild-type C57BL/6 male mice and quantified collagen over protein levels (Fig. 1d-i). We found that the brain and liver had the least amount of collagen to protein levels (0.1%; Fig. 1d), heart and kidney around 1%, muscle and lung around 3.5%, colon 5.7%, skin 20%, bone 25-35%, and tendon about 40% of total collagen to the protein of wild-type CD1 female mice (Fig. 1d; Supplementary Table 3). To provide a reference point, we quantified collagen or protein weight per weight of wet tissue (Fig. 1e, 1f; Supplementary Table 3). Next, we quantified collagen over protein levels of 41 weeks old wild-type C57BL/6 male mice and found higher collagen levels for skin (38.5%) and tendon (50.5%) (Fig. 1g; Supplementary Table 4). This difference could be either due to age, gender, or genotype. Furthermore, we noticed a heterogeneity of collagen to protein levels for individual animals’ organs within the same cohort (dots in the bar graph represent individual mice; Fig. 1d, 1e). For comparison, we searched the literature for collagen measurements normalized to protein levels (BSA, amino acids, nitrogen; Fig. 1j; Supplementary Table 5). We only found a few studies and included rat tissues as additional reference points. Reassuringly, the collagen over protein values were in similar ranges. Thus, our measurements provide reference values for collagen over protein levels for several organs of wild-type mice.

### Quantifying total collagen levels normalized to total proteins of entire mice

An accumulating body of literature states the quote: “30% of total protein is collagen in mammals” and the quote: “collagen is the most abundant protein”; however, a reference to the original work that quantified these percentages is missing (Fig. 2a, Supplementary Table 6). None of the over 50 publications we found had an original reference for these statements (Supplementary Table 6). Our literature search to identify the original work for these percentages led to a review paper from 1961 as a probable source ^35^. In this review article, R.D. Harkness states: “A complete analysis of a whole animal has only been performed on mice (Harkness, Harkness & James, 1958) ^36^. Collagen formed about a fifth of the total body protein.” ^35^. In that referenced study ^36^, Harkness and colleagues dissected nine adult male albino mice, each into five parts (skin, carcass, viscera, femora, quadriceps). They quantified the collagen content of these tissues by multiplying hydroxyproline levels by 7.46 and normalized them to total nitrogen (N) levels as a proxy for protein levels ^36^. Then they added the values of these five parts to estimate the collagen levels of the entire mice. They found that: “Collagen formed 2.6 ± 0.1 % of the body weight and 17.7 ± 0.7% of total N in the control animals” ^36^.

**Fig. 2.**
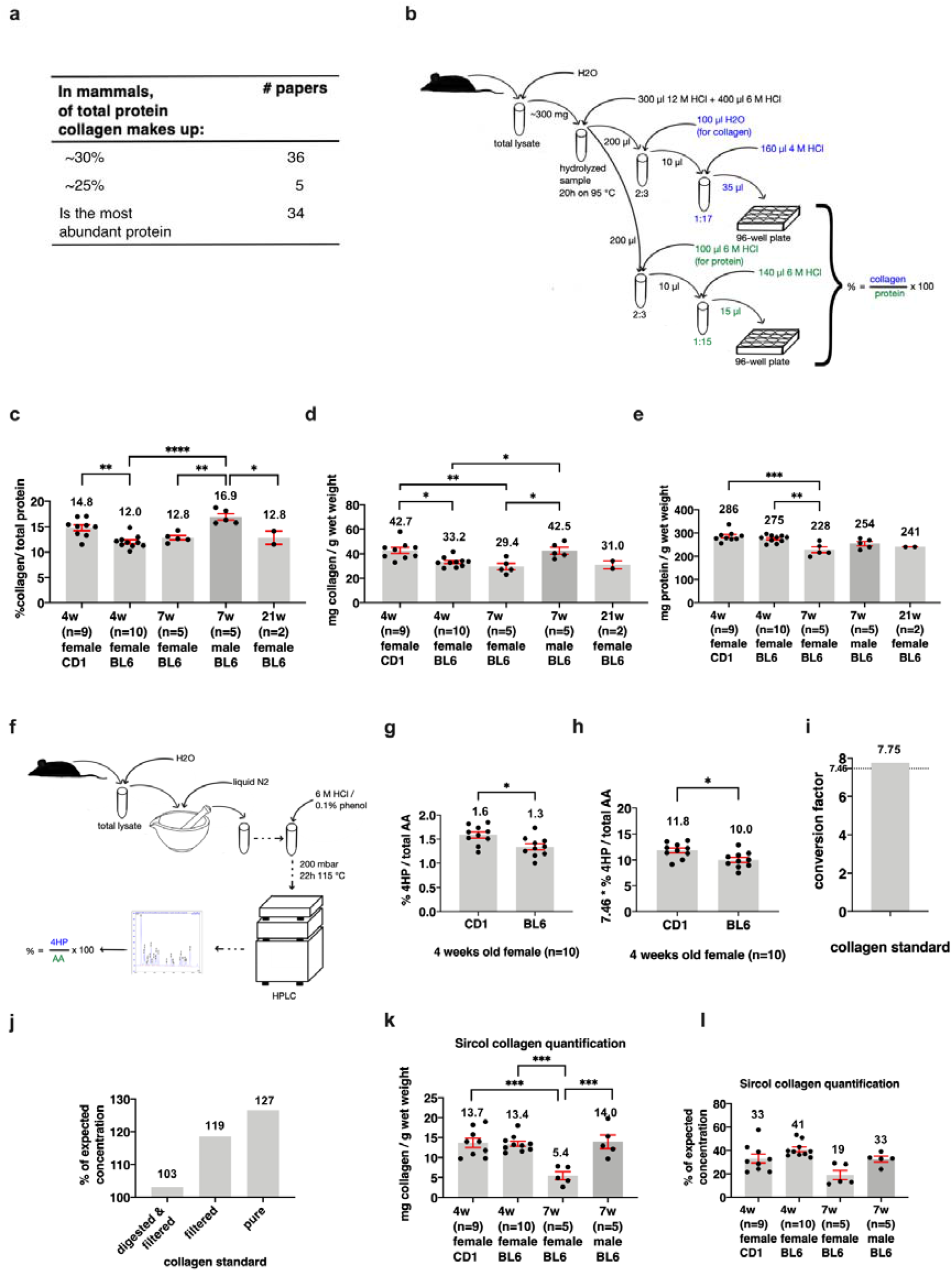
Collagen content of entire mice. **a**, literature indicating a proportion of collagen in mammalian protein. See Supplementary Table 6 for details. **b**, schematic representation of the total collagen and total protein assay procedure. **c**, the percentage of collagen normalized to the total protein of a whole female mouse, calculated as shown in (a). The first bar depicts a CD1 mouse, whereas the others represent C57BL/6 mice. Numbers show the age in weeks (w) or sample size (n). **d**, illustrates the amount of collagen per gram wet weight of a mouse (mg/g). e, shows the portion of protein per gram wet weight (mg/g). c-e, see Supplementary Table 7 for details. f, illustrates the amino acid analysis procedure. Some steps are omitted (dashed lines). For detailed information, see Materials and Methods. g, shows the proportion (%) of 4HP over total amino acids (AA) of the four weeks old female CD1 and C57BL/6 mice. h, depicts the values shown in (f), multiplied with the 4HP to collagen conversion factor (7.46). g and h, see Supplementary Table 8 for details. c-e and g-h, mean ± SEM, shown is the mean of 2-10 biological replicates. Values of biological replicates were derived from 2 (g and h) 3 (c-e) technical replicates. One way ANOVA post-hoc Tukey, **: p≤ 0.01. i, illustrates the experimentally derived conversion factor of the collagen standard. See Supplementary Table 8. j, % of the expected collagen standard concentration used in the previous assays when measured with the Sircol assay (1200 μg/mL equals 100 percent). See Supplementary Table 10. k, illustrates the amount of collagen per gram wet weight (mg/g). See Supplementary Table 10. l, shows the percentage of the expected collagen concentration (see (d)) when measured with the Sircol collagen quantification. See Supplementary Table 10.

There are limitations to this approach of combining and estimating total collagen levels from individual tissue parts. Moreover, quantification of the total collagen of an entire mouse has not been performed before. To address this, we homogenized individual entire mice and hydrolyzed aliquots of these whole mice homogenates with hydrogen chloride to the amino acid levels to determine total collagen over total protein levels of entire mice (Fig. 2b, Materials and Methods for details). We found that 4-week-old outbred wild-type CD1 female mice contained 14.8% of collagen over total protein levels, inbred wild-type C57BL/6 female mice at the age of 4 weeks contained 12.0%, 7- or 21-weeks old had 12.8% of total collagen over total protein (Fig. 2c-e, Supplementary Table 7). Although in wild-type C57BL/6 female mice from 4 to 21 weeks, there was no significant change in collagen over protein levels, 7-week C57BL/6 male mice showed higher collagen over protein levels compared to corresponding 7-week C57BL/6 female mice, 16.9% versus 12.8%, respectively (Fig. 2c, Supplementary Table 7). This suggests a gender-specific effect on total collagen over protein levels. Furthermore, the 16.9% collagen to the protein content of our 7-week C57BL/6 male mice is comparable to the 17.7 % of the male albino mice determined by Harkness and colleagues ^36^.

To complement our colorimetric measurements of hydroxyproline and total amino acids levels, we analyzed amino acid levels within the samples with high-performance liquid chromatography (HPLC; Fig. 2f). We found that hydroxyproline (4HP) over total amino acids is 1.6% and 1.3% of 4-week old wild-type CD1 and C57BL/6 female mice, respectively (Fig. 2g, Supplementary Table 8). To convert the hydroxyproline levels to collagen levels, we used the conversion factor of 7.46. We found 11.8% and 10.0% of 4-week-old wild-type CD1 and C57BL/6 female mice, respectively (Fig. 2h, Supplementary Table 8). The conversion factor of 7.46 is an estimate based on hydroxyproline levels of collagens from various tissues of cows, pigs, sheep, chickens, kangaroos, and rats resulting in 13.0-14.4% of hydroxyproline in mammalian collagens ^32,37^. Since experimental quantification of this conversion factor for mice is missing, we assessed and compared this conversion factor from our data. We found that a conversion factor of 7.46 reflects well collagen estimation from hydroxyproline in whole mouse samples (Supplementary Fig. 2 g-i; Supplementary Table 9). Furthermore, in our HPLC, we also ran a collagen standard to know how much hydroxyproline levels correspond to total collagen levels. When normalizing with this collagen standard, we found 7.75 x (Fig. 2i, Supplementary Table 8). Thus, we obtained comparable results either using a colorimetric approach or HPLC. Taken together, our measurements suggest that, on average female wild-type mice have about 12.3 ± 1.5% of collagen over total protein. Consistent with the previous measurements by Harkness, Harkness & James, 1958 ^36^, male wild-type mice showed about 17% of collagen over total protein.

To assess total collagen levels differently than by using hydroxyproline as a read-out for collagen, we digested our samples with pepsin. We used an adapted version of Sircol staining to quantify soluble collagens ^28^. For Sircol staining, the dye Sirius red binds collagens and other non-collagenous proteins. By running the samples over columns that retain larger proteins, such as collagens, the accuracy for Sircol collagen quantification is improved ^28^. We validated this approach using a collagen standard and found similar collagen content with the Sircol method compared to when assessed by hydroxyproline measurements for pure protein samples (Fig. 2j, Supplementary Table 10). From our whole mouse lysates digested with pepsin, we recovered about 5-14 mg collagen per g wet weight, which is about 19-41% of estimated collagen by hydroxyproline measurement (Fig. 2k and Fig. 2l, Supplementary Table 10). Normalizing the Sircol-derived collagen levels (mg collagen / g wet weight) to our total protein estimates (mg protein / g wet weight), resulted in on average 5% of collagen over total protein (Supplementary Table 10). This estimation is lower compared to the 12-17% of collagen content by the hydroxyproline measurements. We speculate that the underestimation is in part due to the lack of proper solubilization of collagen using this approach.

### Total protein content and amino acid composition of mice

Next, we asked how much total collagen and protein is found in entire mice. In the literature, we found only one publication from 1977 that estimated a total protein of 1.5 g and 1.4 g per 3-week-old male or female CB57BL/6J ob/ob mice, respectively, based on nitrogen levels ^38^. We quantified higher protein levels of 3.8 g of total protein per 4-week-old BL6 wild-type mice (Fig. 3a). We found wild-type mice contain about 4-8 g of total protein and 0.5-1.2 g of total collagen per mouse (Fig. 3a, 3b), which corresponds to 265 mg of protein and 37 mg of collagen per g body weight on average (Supplementary Table 7). The total protein and collagen per mouse were poorly correlated with its total body wet weight (Fig. 3c-f). The collagen levels correlated better, which makes sense since collagens are structural and scaffold proteins important for tissue geometry and size (Fig. 3d).

**Fig. 3.**
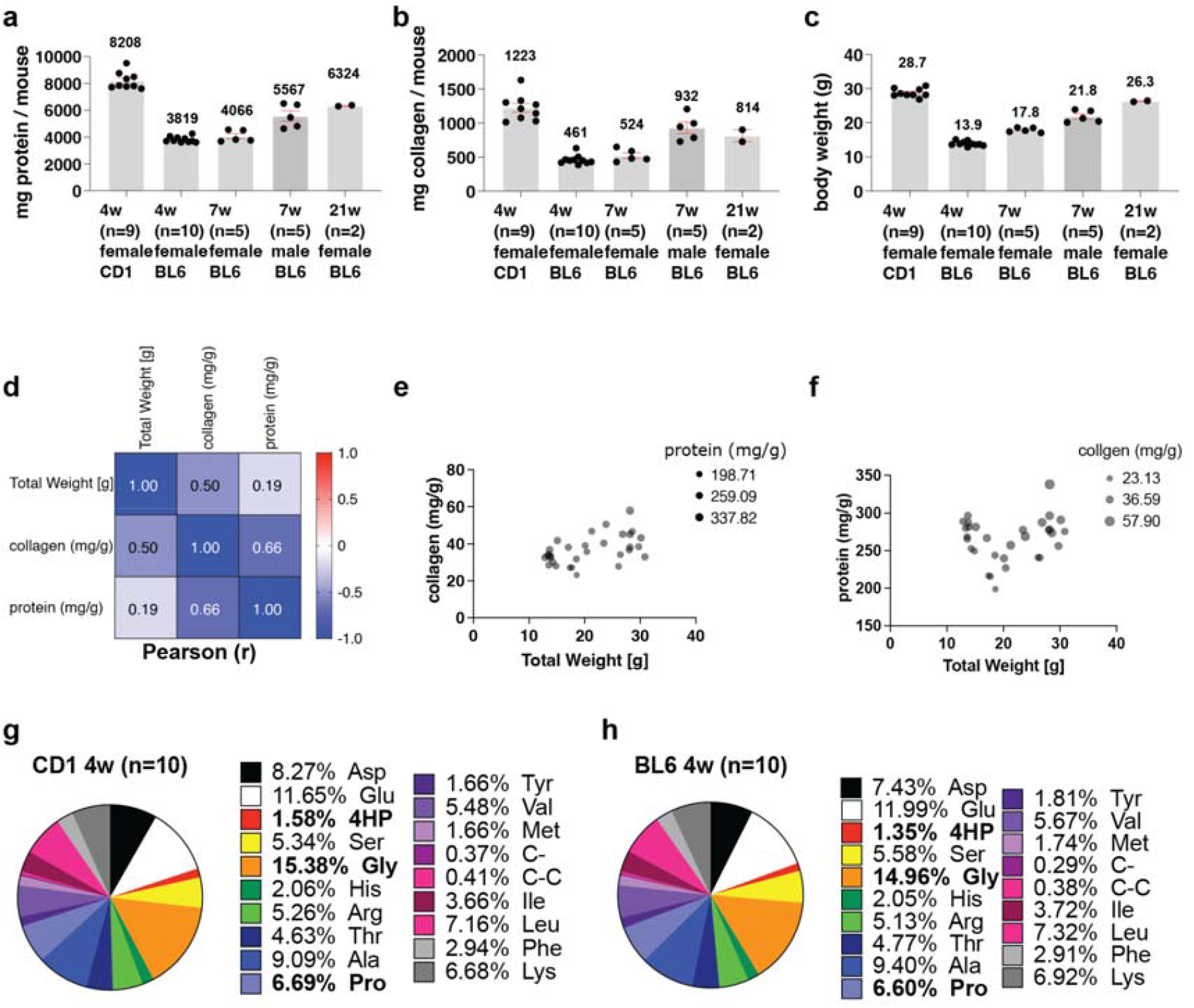
Protein and collagen content of a mouse and its mean body weight. **a**, shows the protein content (mg) of mice. **b**, illustrates the collagen content(mg) of mice. **c**, depicts the mice’s body weight (g). **a-c**, mean ± SEM, shown is the mean of 2-10 biological replicates. Values of biological replicates were derived from 3 technical replicates. **d**, Pearson multivariate correlation matrix. **e**, collagen correlation with body weight (bubble size indicates corresponding protein levels). **f**, protein correlation with body weight (bubble size indicates corresponding collagen levels). See Supplementary Table 7 for details. **g**, Pie chart representation of the amino acid composition of 4 weeks old CD1 mice (n= sample size). See Supplementary Table 8. **h**, Pie chart representation of the amino acid composition of 4 weeks old C57BL/6 mice. See Supplementary Table 8.

Next, we asked what is the relative amino acid composition of mice. We found that glycine with 15% was the most abundant amino acid in mice (Fig. 3g-h, Supplementary Table 8). Proline was 6.5%, and hydroxyproline (4HP) was about 1.5% relative to all other amino acids (Fig. 3g-h, Supplementary Table 8). Thus, we provide the first relative amino acid composition of entire mice.

### Collagen quantification using proteomics

Next, we used a quantitative proteomics approach. To enrich extracellular matrix proteins, we separated whole mouse lysates by SDS-PAGE, gel lanes were cut into five slices per sample, and proteins therein were digested with trypsin overnight. Generated peptides were extracted and analyzed by LC-MS/MS (see Methods for details). Extracted ion currents of peptides were used for label-free quantification employing the iBAQ algorithm ^39^. For each sample, the relative proportion of all quantified collagen proteins was calculated in comparison to the overall intensity of all proteins. We quantified 4.6±0.3% for 4-week-old CD1 mice, 4.6±1.4% for 4-week-old C57BL/6 mice, and 6.4±0.4% for 21-week-old C57BL/6 mice (Supplementary Table 11). Although these percentages are similar to the Sircol collagen quantifications, these percentages were lower compared to the hydroxyproline measurement, probably due to the crosslinked nature of collagens, and we would expect that trypsin is less efficient in generating measurable peptides.

### Matreotype of entire mice

To establish matreotypes ^15^, which is the composition of the ECM (*i*.*e*., matrisome) associated with entire mice, we analyzed our ECM-enriched proteomics data according to matrisome category ^40^. We detected 167-233 of the 1110 mouse matrisome ^41^ proteins in 4-week-old CD1, 4-week-old C57BL/6, and 21-week-old C57BL/6 wild-type mice (Fig. 4, Supplementary Table 11). To our surprise, serine protease inhibitors (Serpina1,3) were more abundant than collagen type I (Col1a1,2), followed by fibrillin (Fbn1), elastin (Eln), and collagen type 6 (Col6a1,2) (Supplementary Fig. 3, Supplementary Table 11). Furthermore, considering all detected proteins, albumin (Alb), hemoglobin (Hbb), and parvalbumin (Pvalb) were the most abundant protein, followed by histones, actin (Acta1), serine protease inhibitors (Serpina1,3), and then collagen type I (Col1a1,2) (Supplementary Fig. 3, Supplementary Table 11). Thus, we provide the matrisome atlas of entire mice and found that collagens are not the most abundant protein.

**Fig. 4.**
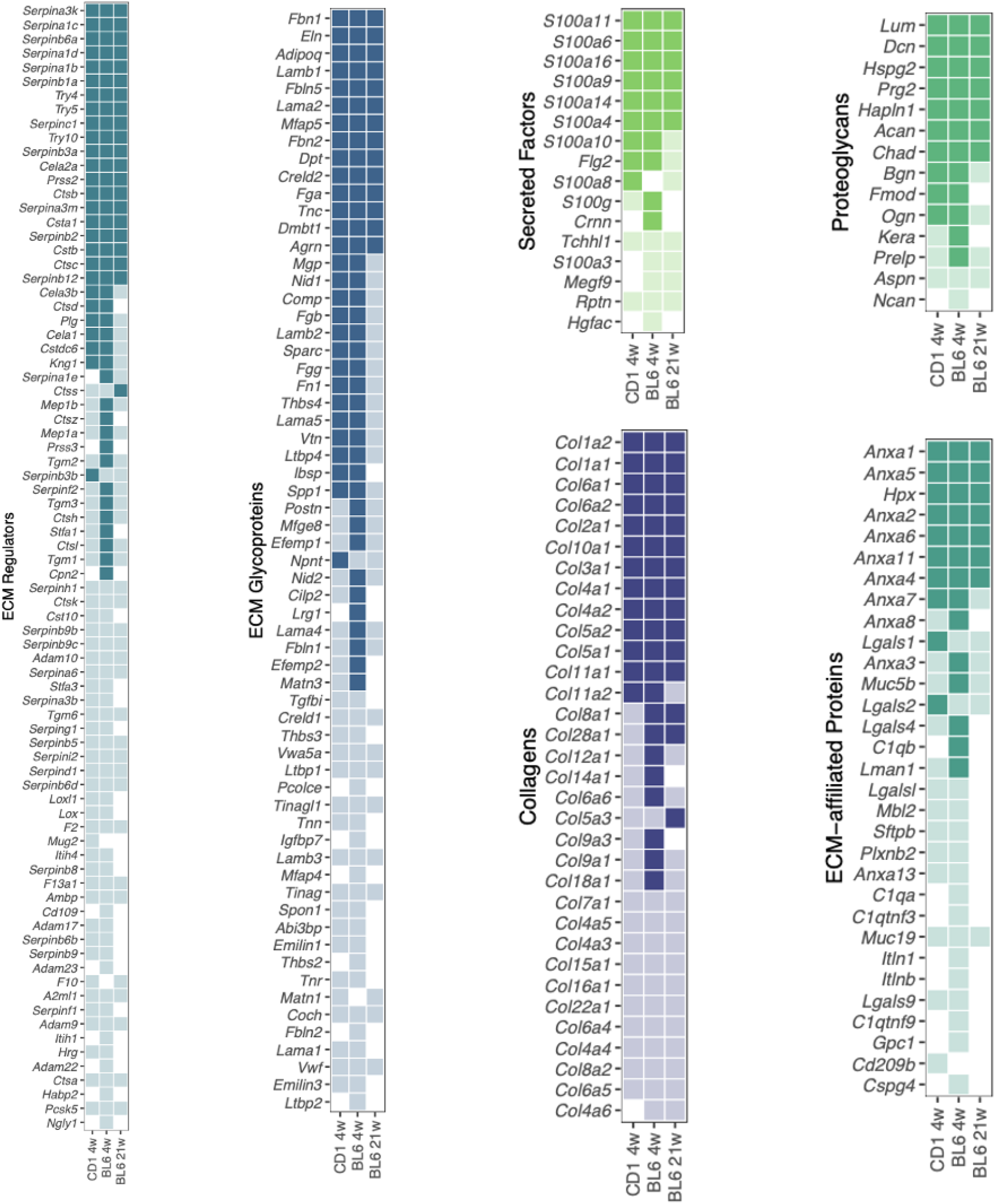
Matreotpype atlas of the wild-type murine proteome. Comparative analysis of all matrisome proteins which could be quantified in at least one of the three animal cohorts: wild-type CD1 (4 weeks) and **C57BL/6** at four and twenty-one weeks old. Solid squares represent proteins that were observed to be abundant (> median) in the respective cohort, transparent squares indicate low abundance (<= median), and white squares highlight non-observable protein-cohort combinations. Details are in Supplementary Table 11.

## Discussion

Biomedical engineering and science rely on previously quantified reference points to build models and compare health versus disease states. Collagen has been assumed to be the most abundant protein of mammals making up two-thirds of total protein. However, over 50 studies ranging from material engineering to cancer research relied on these reference points without an original reference of such quantification.

Although nowhere directly stated, this statement might be based on a review paper from 1961 stating that “collagen formed about a fifth of the total body protein” ^35^. This statement was based on the authors’ previous study from 1958, where nine male mice carcasses were divided into five parts, and hydroxyproline levels over total nitrogen levels were determined, which were added together, resulting in 17.7% of total collagen over protein ^36^. Astonishingly, our quantification revealed a similar percentage of 16.8% of total collagen over protein of entire male mice. This suggests that we had a correct reference point but, for an unknown reason, was inflated to 30%, an interesting hyperbole that might be worth examining by science philosophers.

Besides this, we find sex-specific differences in collagen over protein for females of 12% and males 17%. We provide reference points for collagen over protein levels for ten tissues. Using proteomics, we find that collagen is not the most abundant protein. We provide the matreotype of entire mice as a reference point for an ECM protein atlas. In addition, we provide reference points for milligram protein per gram mice and amino acid compositions, whereby glycine is the most abundant amino acid.

Although not systematically examined, there were few reports of collagen over protein ratios for mice and rat tissues. Our estimates were similar to previous estimates even across species. Also, estimating collagen over protein levels using colorimetric compared to HPLC of hydroxyproline levels was similar. However, using collagen staining and proteomics approaches showed internally consistent collagen percentages of 5-6%, which are lower than when quantified the same animals with hydroxyproline quantifications. For both methods, samples need to be separated via either running on a gel or on a column, probably losing material. Furthermore, a lower yield might also be due to the inherent insolubility of the collagen networks ^42^, and in the case of proteomics, limited protease access to crosslinked peptides and lack of identification of crosslinked peptides, limitations we had not addressed here. A previous study on tendons and ligaments achieved complete solubilization by successive protease and elastase digestion, which allowed the harvesting of more collagen and elastin ^43^. Thus, such an approach might allow quantifying more collagen over protein levels, which might asymptote to our hydroxyproline to collagen estimations.

Our proteomics analysis revealed collagen type I (Col1a1,2), fibrillin (Fbn1), elastin (Eln), and collagen type 6 (Col6a1,2) as the most abundant core matrisome proteins. This is comparable to a recent meta-analysis of different proteomics studies on articular cartilage, bone, ligament, skeletal muscle, and tendon ^44^. It is also similar to tissues with less ECM, such as mammary glands and liver ^45,46^. As in our study of entire mouse proteomics, collagen type I is the most abundant collagen in all tissues but shows a wide range across different tissues. This suggests that collagen type I is the most abundant collagen in mammals across tissues.

In summary, we quantified collagen levels for mice tissues and entire mice. We reconcile assumptions made in textbooks and research papers. Thus, we provide several important reference points to help interpret new findings related to collagen levels.

## Methods and Materials

### Sample preparation

#### Mouse Tissue

CD1 mice were provided by the institute of laboratory animal science (University of Zürich; AniMatch). The Veterinary office approved all other animals of the Canton of Zürich (Licence Nr. ZH092-19). All experiments were performed post-mortem.

#### Total mouse lysate

We used inbred C57BL/6 and outbred CD1 mice for the total lysate experiments (9 to 10 CD1 and 10 C57BL/6 mice, all 4 weeks old female, and 12 C57BL/6 mice (male: 5 × 7 weeks, female: 5 × 7 weeks and 2 × 21 weeks). These mice were euthanized, frozen at -20°C and then individually mixed with a NutriBullet 600 Series after having added 40 mL of distilled water (Sigma-Aldrich) to the 4 weeks old C57BL/6 mice, 50 mL to the CD1, and 20 mL to the 7-21 weeks old C57BL/6 mice. This mixture was filled into 50 mL tubes and was bead-bounced for 3 × 10 to 6 × 10 minutes. Then 500 μL was aliquoted into cryotubes and sonicated for 20 seconds to homogenize it completely. Despite homogenizing, the hair remained in the samples and was not filtered out. The aliquots were snap-frozen and stored at -80°C.

#### Organ samples

Mice were euthanized, and skin punches (5 mm, Biopsy punch, Stiefel) were collected from the shaved back. To harvest the tail skin, the tail was cut, and the skin was flayed. A 5 mm piece of the tail skin was cut with a razor blade 3 cm and 3.5 cm away from the tip because, at this distance, the skin looked uniformly in the different age cohorts. Lung, heart, liver, colon, kidney, skull, brain, femur, tail, and hamstring muscle were dissected, snap frozen, and stored at -80°C.

### Amino acid analysis

Whole mouse lysate aliquots were pestled in liquid nitrogen and sent to Analytical Research Services (Bern, Switzerland) for amino acid analysis. They measured the probes according to Bidlingmeyer *et al*. ^*47*^. In brief, 25 μL of the homogenate was pipetted for hydrolysis into a glass tube (6 × 50 mm) and vacuum-speed dried, then 25 μL 6M HCl with 0.1% phenol was added. For hydrolysis, 200 μL 6M HCl / 0.1% phenol was pipetted into a vessel with frit, and the glass tube was placed inside. The vessel was flooded with nitrogen and sealed under a vacuum (ca. 200 mbar). Samples were hydrolyzed for 22h at 115°C. Then they were dried and resuspended in 350 μL 0.1% trifluoroacetic acid (TFA). After a short centrifugation, the supernatant was diluted 1:10 with water. An aliquot of 10 μL was transferred into a tube and dried. After the reaction with phenylisothiocyanate (PITC) the aliquot was dried again and dissolved in 50 μL high-performance liquid chromatography (HPLC) Eluent A. 20 μL was injected into an HPLC (Dionex with P680 HPLC pump, ASI 100 (automated sample injector), thermostatted column compartment TCC-100 and ultimate 3000 RS diode array detector).

### Quantification of total collagen and protein

#### Total mouse lysates Collagen and protein assay

To quantify the total collagen normalized to the total protein of the total lysate, the QuickZyme total collagen and total protein assays (Biosciences) were performed according to the manufacturer’s protocol. The QuickZyme total collagen assay is based on a colorimetric read-out for free hydroxyproline amino acids. Free hydroxyproline is oxidized with Chloramine-T to pyrrole and then stained with Ehrlich’s reagent ^48^. Pure rat tail collagen was hydrolyzed, and different concentrations were loaded according to the protocol on the assay plate to be able to generate a standard collagen curve, which was used to determine the total collagen content in our samples.

The QuickZyme total protein assay is based on a colorimetric read-out. Hydrolyzed free amino acids are assessed by using genipin, and the color change acts as a read-out ^34^. According to the protocol, a standard curve of pre-hydrolyzed bovine serum albumin (BSA) was employed to determine total protein concentration. Before conducting the assays, the samples were vortexed for 10 seconds. 300 mg of the aliquot was used for the lysis, ensuring that all hair in the sample was weighed. 300 μL 12M HCl (VWR chemicals) and 400 μL 6M HCl were added. This mixture was hydrolyzed at 95°C for 20h to the amino acids. The hydrolyzed samples were centrifuged for 10 min at 13000 x g. The amino acid homogenate was split to assess collagen and protein levels.

For the collagen assay, the hydrolysate was diluted (2:3) with water to reach a final HCl concentration of 4M. The same dilution was conducted with the samples for the protein assay but with 6M HCl. Then the samples were further diluted to be in the range of the standard curve (collagen samples dilution was 1:17 with 4M HCl, and protein samples dilution was 1:15 with 6M HCl). These dilutions were loaded on the 96-well plates (one for the collagen assay and one to conduct the protein assay), and the buffer and color reagent required were added. After 1h incubation time at 60°C for the collagen plate and at 85°C for the protein plate, the plates were placed on ice for 5 min or 7 min, respectively, and absorbance was measured at 570 nm (CLARIOstar, BMG Labtech, CLARIOstar-Data Analysis, 3.1).

To calculate the total milligram collagen or protein per gram of an entire mouse, the dilution factor for homogenizing the mouse was considered.

#### Hydroxyproline assay

Hydroxyproline concentrations were measured according to the Hydroxyproline Assay Kit (Sigma-Aldrich) protocol. This assay also detects free 4HP that is oxidized with Chloramine-T to pyrrole and stained with Ehrlich’s reagent ^48^. Samples were diluted 1:2 with 12 M HCl and hydrolyzed at 120°C for 3 hours. Activated charcoal was added, and tubes were centrifuged at 10000 x g for 3 minutes. 50 μL of the standard collagen samples (see 3.2.3.1.3) were loaded onto the plate, and the other samples were diluted further in two steps (1:10 and 1:9) to be in the range of the hydroxyproline standard curve. Into the plate, 50 μL of the final dilution was loaded and dried at 60°C. 100 μL of Chloramine T/Oxidation Buffer mixture was added and incubated at room temperature for 5 minutes. 100 μL of the diluted DMAB reagent (DMAB: perchloric acid/Isopropanol=1:1) was added to each well and incubated for 90 minutes at 60°C. Absorbance was measured at 560 nm. The results were multiplied by the hydroxyproline to collagen conversion factor (7.46), as stated in Neuman *et al*. ^*49*^.

#### Collagen and hydroxyproline standard preparation

To verify the collagen content of the samples, standard collagen concentrations, and hydroxyproline standard concentrations were loaded on the assay plates used for the assays comparison and prepared as follows.

For the collagen assay, the collagen assay standard (rat tail collagen, 1200 μg/mL (Bioscience)) was prepared according to protocol. To be able to generate a 4HP standard curve on the collagen assay plate, the 4HP standard (1 mg/mL (Sigma-Aldrich)) was diluted 1:10 with HCl (0.1 mg/mL in 6M HCl). 125 μL of this concentration was mixed with 62.5 μL 4M HCl and 62.5 μL water. 80 μL of this dilution was transferred and mixed with 20 μL 4M HCl (40 μg/mL). The further standard concentrations were reached by consecutive 1:2 dilutions. (20 μg/mL, 10 μg/mL, 5 μg/mL, 2.5 μg/mL). 4M HCl was used as blank.

For the 4HP assay, 250 μL of the collagen standard was diluted with 250 μL water (600 μg/mL). This solution was diluted 1:2 with water (300 μg/mL). 240 μL of the last dilution was mixed with 120 μL of water (200 μg/mL). A 180 μL aliquot was transferred to a new tube and diluted with 180 μL H2O (100 μg/mL). This 1:2 dilution was repeated 4 times (50 μg/mL, 25 μg/mL, 12.5 μg/mL, 6.25 μg/mL), and a water blank was prepared. To be in the range of the 4HP standard curve, 150 μL of each sample and blank were pipetted into a new tube and diluted with 150 μL 12M HCl.

After hydrolyzation (3h at 120°C), 50 μL of each sample was pipetted on a 96-well plate. The hydroxyproline standard (1 mg/mL) was prepared according to the protocol for the 4HP assay.

#### Sircol™ Soluble Collagen Assay

Total collagen concentrations were also measured with the Sircol™ Soluble Collagen Assay (biocolor life science assay). 100 mg of the total lysates were mixed with 1mL 0.1 mg/mL pepsin in 0.5M acetic acid (Merck) and put at 4°C overnight. The next day, 500 μL 0.32 mg/mL pepsin in 0.5M acetic acid (Merck) was added and again left overnight at 4°C. 1 mL of this solution was then transferred into an Amicon Ultra-2 Centrifugal Filter Unit (Millipore). This unit was centrifuged for 60 minutes at 3220 rcf. The flow-through was discarded, and the collagen remaining in the filter was collected by spinning the inverted tube for 2 minutes at 1000 rcf. 0.5M acetic acid (Merck) was added to reach 500 μL. 50 μL of this solution was pipetted into a new tube, and the Sircol assay was performed according to protocol. Finally, 1 mL alkali reagent was added. 200 μL was pipetted on a 96-well plate.

The rat tail collagen standard, 1200 μg/mL (Bioscience) used in the collagen assay was also measured with three different pre-treatments. In the first treatment, 100 μL of the standard was digested and filtered like the samples described above. 150 μL 0.5M acetic acid was then added to the collagen collected in the tube. For the second, 120 μL of the standard was diluted with 880 μL 0.5M acetic acid and only filtered like the samples. This time 0.5M acetic acid was added to reach 200 μL. 50 μL of each collagen standard sample was transferred in separate tubes, and 50 μL 0.5M acetic acid was added. Finally, 30 μL of the untreated collagen standard was diluted with 70 μL 0.5M acetic acid. The Sircol assay was performed according to protocol. After having added 1 mL alkali reagent, 200 μL of the differently treated standards were transferred to a 96-well plate.

#### Bicinchoninic acid (BCA) protein assay

Total protein concentrations were also measured with the Pierce™ BCA Protein Assay Kit (Thermo Scientific). This assay combines the protein-induced biuret reaction with the colorimetric detection of the resulting cuprous cation (Cu^1+^) by BCA ^50^.

The standard for this assay was prepared according to the protocol. In the first replicate, samples were vortexed, and hairs were included. For the second replicate, the samples were sonicated for 20 seconds, and hairs were excluded, and for the last replicate, samples were not pretreated. The total lysate was diluted (3:10 and 2:3). 25 μL of the final dilution was pipetted into a 96-well microplate. 200 μL of the BCA working reagent was added before shaking the plate for 30 seconds on a plate shaker. Afterward, the plate was covered and incubated at 37°C for 30 minutes. The plate was cooled to room temperature, and absorbance was measured at 562 nm.

#### Organs

To quantify collagen normalized to protein levels in organs, the QuickZyme total collagen and total protein assays (Biosciences) were conducted. For most organs, 0.1 g of frozen organ tissue was weighted and used for the assay. For skin tissue, a skin punch (or half of a skin punch) was weighted, and for the skull and femur, 0.01 g was used. Organs were diluted with 6M HCl using the following formula: v= aw x D_organ_ -aw (v = μL 6M HCl added to organs, aw = organ assay weight (mg), D_organ_= experimentally determined organ specific dilution factor, D_colon_ = 12.5, D_femur_= 74.2, D_lung_ =10.9, D_skin or tendon_= 73.5, D_muscle_ = 11.2, D_brain_ = 5.7, D_heart_ = 11.0, D_liver_ = 7.3, D_skull_ = 98.2, D_kidney_ = 10.8) and hydrolyzed at 95°C for 20h. The hydrolyzed samples were processed like the total mouse lysates, meaning diluted to 4M HCl (2:3) with water if collagen was intended to be measured. Protein samples were diluted 2:3 with 6M HCl. To finally be within the range of the standard curve, the samples were diluted further. The liver and brain samples were diluted 1:1 for collagen and 1:10 for protein measurements. Tendon samples were processed like skin samples, but collagen samples were diluted 1:16. All other organs were diluted 1:8 for protein and collagen measurements.

### Statistical analysis

Statistical analysis was performed with GraphPad Prism 8.0 using a one-way or two-way analysis of variance (ANOVA) followed by a posthoc Tukey’s, Dunnett, or Sidak multiple comparisons test or a two-tailed unpaired t-test with a 95% confidence interval. Normal distribution of the values was assumed. Values represent means ± SD. * *P* ≤ 0.05, ** *P* ≤ 0.01, *** *P* ≤ 0.001, and **** *P* ≤ 0.0001.

### Quantitative proteomic analysis of mouse samples

Mouse samples were dissolved in 10 mM Tris-HCl buffer containing 4% SDS, pH 7.5, followed by sonication for 2 minutes. SDS-PAGE loading buffer was added, and protein samples were heated for 10 minutes at 95°C, treated with 1 mM DTT, and alkylated using 5.5 mM iodoacetamide for 10 minutes at room temperature. Samples were fractionated on 4-12% gradient gels, and proteins were in-gel digested with trypsin (Promega), five fractions per sample. Tryptic peptides were purified by STAGE tips, and LC-MS/MS measurements were performed on a QExactive Plus mass spectrometry coupled to an EasyLC 1200 nanoflow-HPLC (all Thermo Scientific). MaxQuant software (version 1.6.2.10) ^51^ was used for analyzing the MS raw files for peak detection, peptide quantification, and identification using a Uniprot mouse database (version April 2016). Carbamidomethylcysteine was set as fixed modification, and oxidation of methionine was set as variable modification. The MS/MS tolerance was set to 20 ppm, and four missed cleavages were allowed for Trypsin/P as enzyme specificity. Based on a forward-reverse database, protein and peptide FDRs were set to 0.01, the minimum peptide length was set to seven, and at least one unique peptide had to be identified. The match-between run option was set to 0.7 minutes. MaxQuant results were analyzed using Perseus software (version 1.6.2.3) ^52^.

To calculate the relative abundance of collagen proteins compared to the detected proteome, replicate iBAQ values of respective groups were averaged, and the percentage of collagens was obtained by this formula: sum of collagen iBAQ values*100/sum of iBAQ values of all proteins.

### Matrisome annotation of proteomics data

Peptide annotations below a Q-value of 0.05 were augmented with the murine matrisome annotations (http://matrisomedb.pepchem.org) ^40^. Protein abundances for each mouse strain and age group are illustrated by the matrisome category as a tile map. Mean abundances above the overall median protein abundance are displayed in full color, transparent if below, and white if not detected.

## Figures

**Supplementary Fig. 1.**
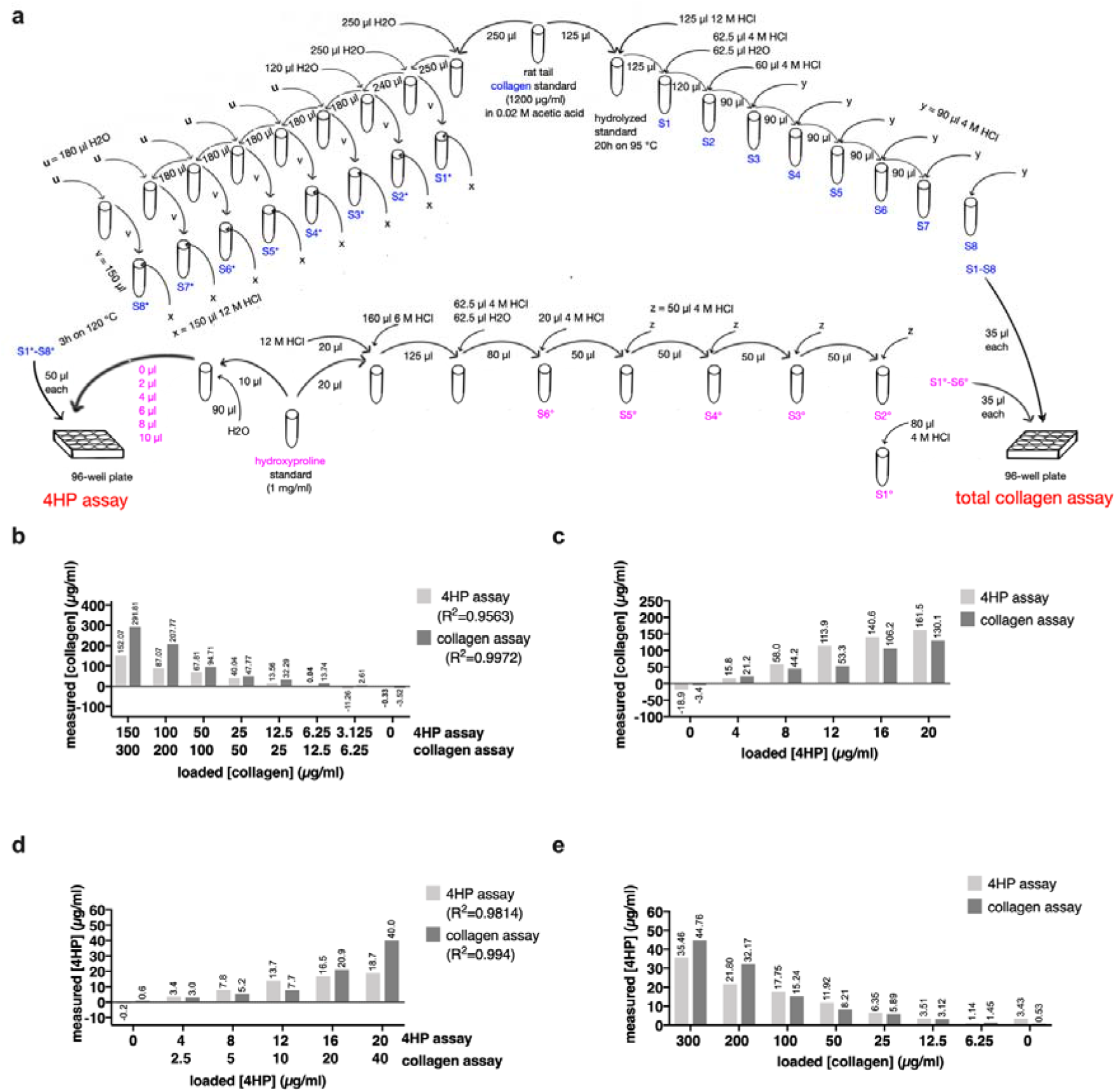
Collagen validation standard curves of the 4HP and collagen assay. **a**, schematic representation of the 4HP and total collagen assay standard curve preparation. The collagen assay standard (rat tail collagen, 1200μg/mL) was prepared according to the protocol for the collagen assay (right). For the 4HP assay (left), 250μL of the collagen standard was diluted with 250μL water (600μg/mL). This solution was diluted 1:2 with water (H2O) (300μg/mL). 240μL of the new dilution was mixed with 120μL water (200μg/mL). A 180μL aliquot was transferred to a new tube and diluted with 180μL H2O (u) (100μg/mL). This 1:2 dilution was repeated 4 times (50μg/mL, 25μg/mL, 12.5μg/mL, 6.25μg/mL). S1* is the blank. To be in the range of the 4HP standard curve, 150μL (v) of each sample was pipetted into a new tube (S1*-S8*) and diluted with 150μL 12M HCl (x). After hydrolyzation (3h on 120°C), 50μL of each sample was pipetted on a 96-well plate. The hydroxyproline standard (1mg/mL) was prepared according to the protocol for the 4HP assay (left). The 4HP assay concentrations need to be divided by 0.05 to convert the concentrations from μg/well into μg/mL (since 50μL of each S* were loaded). To be able to generate a 4HP standard curve on the collagen assay plate (right), the standard was diluted 1:10 with HCl (0.1mg/mL in 6M HCl). 125μL of this concentration was mixed with 62.5μL 4M HCl and 62.5μL water. 80μL of this dilution was transferred and mixed with 20μL 4M HCl (S6°= 40μg/mL). The further standard concentrations were reached by consecutive 1:2 dilutions. (S5°= 20μg/mL, S4°= 10μg/mL, S3°= 5μg/mL, S2°= 2.5μg/mL). S1° is the blank. With the concentrations mentioned, collagen and a 4HP standard curve were generated for each assay. b and d, depict the measured collagen (b) and 4HP (d) values (μg/mL), which were used to generate the standard curve and correspond to the loaded, known concentrations. The top row on the x-axis indicates the concentrations (μg/mL) loaded in the 4HP assay, and the bottom row shows the ones loaded in the collagen assay. R^2^’s represents the coefficient of determination of the resulting standard curves. c, compares the measured collagen values (μg/mL) of the two assays correlating with the loaded 4HP concentrations (μg/mL). For this, the collagen standard curve was considered on the two plates. Since different 4HP concentrations were loaded on the two assays, the rule of proportion was used to make the assays comparable. e, compares the measured 4HP values obtained from the loaded collagen concentrations. This time the 4HP standard curve was used on both plates.

**Supplementary Fig. 2.**
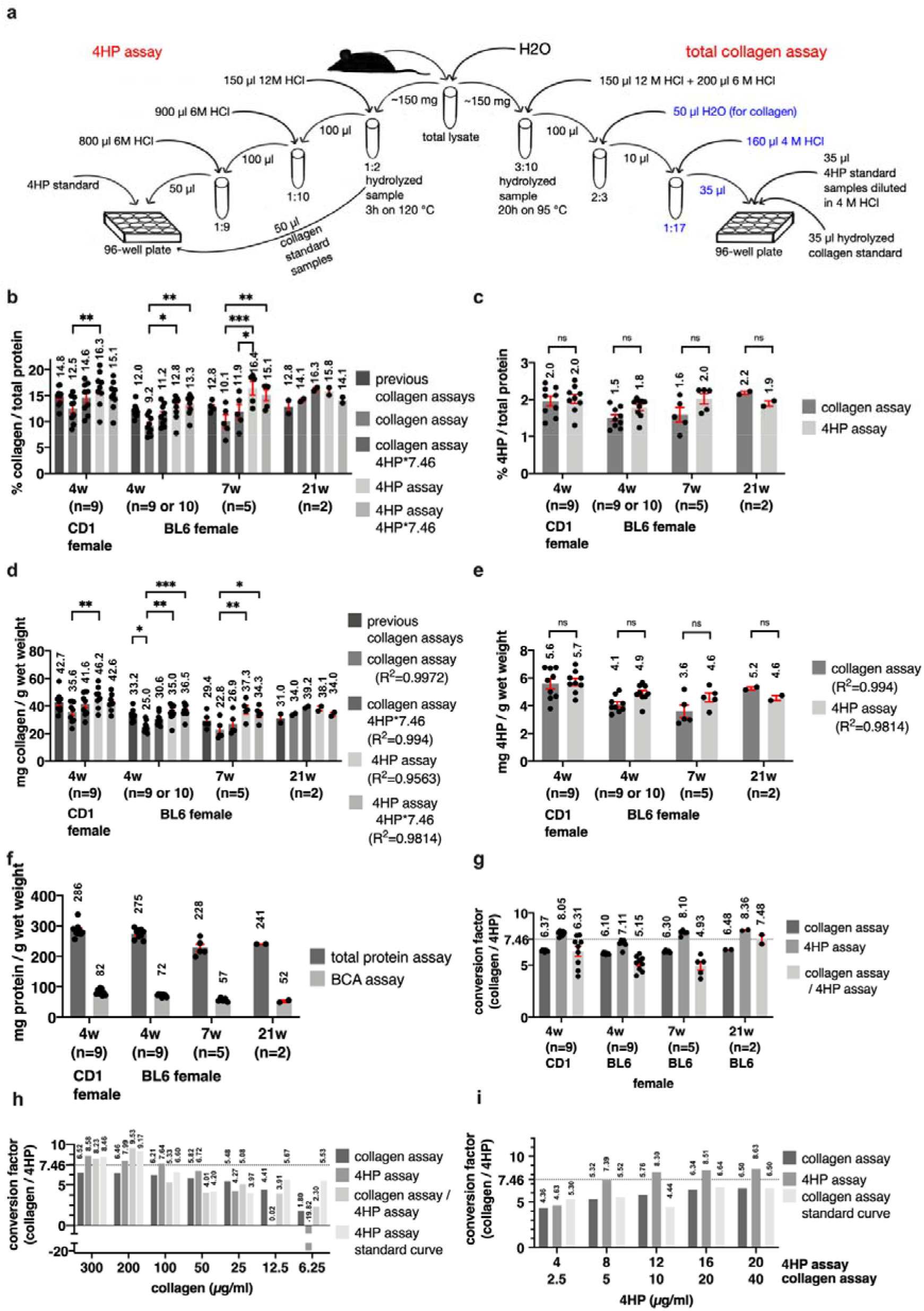
Comparison of the 4HP and collagen assay. **a**, schematic representation of the 4HP and total collagen assay procedure. The mice were diluted with water and aliquoted. An equal amount of the total lysate was used for the 4HP assay and for the collagen assay. For the total collagen assay, the lysates were diluted 3:10 to a final 6M HCl concentration and hydrolyzed for 20h at 95°C. This hydrolysate was diluted 2:3 with water and 1:17 with 4M HCl. 35 μL of the normal hydrolyzed collagen standard, 35 μL of 4HP standard samples derived from the 4HP assay standard solution, and 35 μL of the samples were loaded on a 96-well plate. For the 4HP assay, the lysates were diluted 1:2 with 12M HCl and hydrolyzed for 3h at 120°C. The hydrolysate was further diluted (1:10 and 1:9) with 6M HCl. 50 μL of these samples and 50 μL of the collagen standard dilutions, which were not further diluted after hydrolyzation, were loaded on a 96-well plate next to the normal 4HP standard. **b** and **c**, show a percentage of collagen (b) or 4HP (c) to protein comparison of the different assays. The mean of the 3 previous protein assays was used for normalization. **d** and **e**, compare values (mg/g) of collagen or 4HP, respectively. **b** and **d**, 7.46 is the 4HP to collagen conversion factor cited in the literature (Neuman and Logan, 1950). The 4HP concentration of the corresponding assay is multiplied by this factor to get the collagen concentration. **f**, depicts the protein levels (mg/g) from the total protein assay and the BCA assay. **g**, illustrates the means ± SEM of the empirical conversion factors of the animal groups mentioned on the x-axis. The factor is derived by dividing the collagen concentration of a sample by its 4HP concentration. Three different methods were used to calculate it. Either the values used for the calculation were taken from the same plate (collagen assay, 4HP assay) or from different plates (collagen assay / 4HP assay). **b-g**, the first group consists of 4 weeks old CD1 females. The others are C57BL/6 females. The numbers indicate the age in weeks (w). Shown is the mean ± SEM of 2-10 biological replicates. (values of biological replicates are derived from 1-3 technical replicates), 2way ANOVA posthoc Tukey (a,c) or Sidak (b,d). **h** and **i** show the conversion factors calculated by using the collagen and 4HP standard curves, respectively. Four or three different methods were used to calculate it. Either the values used for the calculation were taken from the same plate (collagen assay, 4HP assay), from different plates (collagen assay / 4HP assay), or the calculation was done by using the measured values and the concentrations indicated in the x-axis (4HP assay standard curve or collagen assay standard curve, respectively). See Supplementary Table 9 for details.

**Supplementary Fig. 3.**
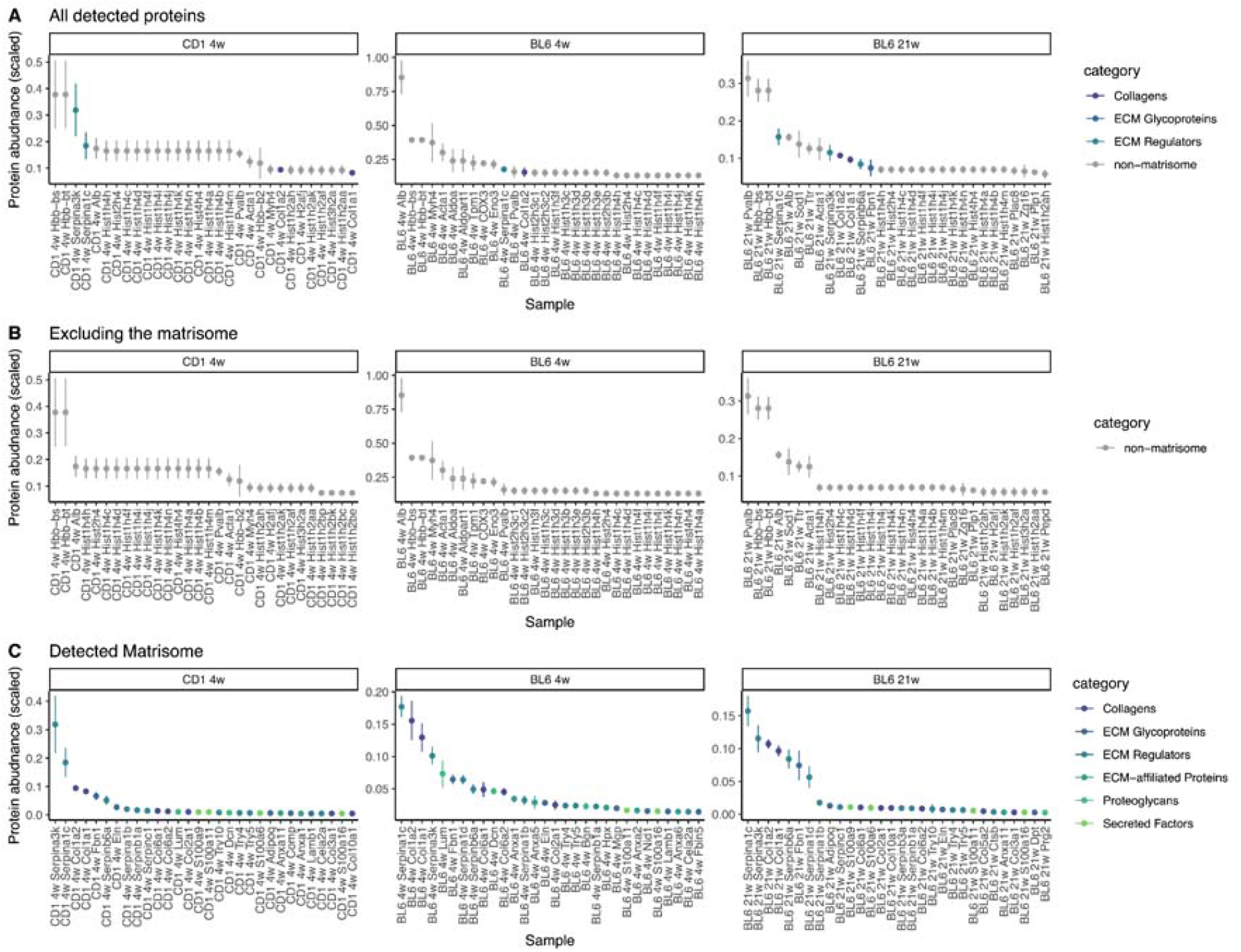
Top 30 most abundant proteins of wild-type mice. The scaled protein abundances are displayed in horizontal panels for each mouse cohort (CD1 4 weeks, BL6 4 weeks, BL6 21 weeks) for all detected proteins (a), all non-matrisome proteins (b), and exclusively the matrisome (c). The matrisome membership is represented by the color of each data point indicating its matrisome category. Details are in Supplementary Table 11.

## Author Contributions

All authors participated in designing the research, executing the experiments, and analyzing and interpreting the data. KT and CYE wrote the manuscript in correspondence with the other authors.

## Acknowledgments

We thank Dr. Cyril Statzer for his assistance in the analysis and visualization of the proteomic dataset, Jasmin Mantl for help harvesting samples and initial work establishing protocols, Philippe Bugnon (LTK1 Kurse) and AniMatch for CD1 mice, Frank Zaucke for literature references, John Hourihan, Jan Gebauer, Mats Paulsson, Matthias Chiquet, Peter Bruckner, and Ewald lab members for discussion and comments on the manuscript.

